# The rise of cheats during experimental evolution is restricted by non-kin interactions between *Bacillus subtilis* soil isolates

**DOI:** 10.1101/2024.03.29.587313

**Authors:** Katarina Belcijan Pandur, Barbara Kraigher, Ana Tomac, Polonca Stefanic, Ines Mandic Mulec

## Abstract

Cooperative behaviors in human, animal, and even microbial societies are vulnerable to exploitation. Kin discrimination (KD) has been hypothesized to help stabilize cooperation. However, the mechanisms that sustain cooperative behavior remain poorly understood. We here investigate the role of KD in limiting the rise of cheats during surfactant dependent cooperative swarming over surfaces by bacterium *Bacillus subtilis* as a model organism. We show that mixing surfactant secreting cooperators and cheats that do not produce surfactants leads to cooperation collapse. However, when such mixed swarms transiently encounter non-kin *B. subtilis* swarms, the frequency of the surfactant nonproducers decreases, suggesting that kinship dependent interactions may limit cheats’ advantage. To further validate this hypothesis, we subjected wild-type co-operators to transient encounters with kin and non-kin swarms over 20 cycles of experimental evolution. Evolved populations exposed to non-kin swarms exhibited lower occurrences of genotypes with defective swarming phenotypes compared to those encountering kin swarms. These results provide compelling support for the prediction that the evolution of cheats in bacterial populations is impeded by kin discrimination providing experimental proof of its role in stabilizing cooperative behavior.

## INTRODUCTION

Bacteria have developed intricate mechanisms to cooperate with each other, often forming complex communities and exhibiting collective behaviors. Although cooperation is widespread among organisms of different organizational levels[1], it is threatened by cheats (also referred to as exploiters) that benefit but do not contribute to the common good [2]. The theory predicts that the evolutionary persistence of cooperative behavior must rely on mechanisms that limit exploiter’s advantage and consequently stabilize cooperative acts [3–5]. For example, cooperation may be more evolutionary stable if individuals preferentially cooperated with highly related (kin) individuals and avoid or antagonize less related (non-kin) conspecifics [5–7]. However, experimental proof revealing non-kin interactions as the cooperation stabilizing mechanism during evolution of bacterial populations is limited.

*Bacillus subtilis*, a gram-positive bacterium, exhibits a rich arsenal of social behaviors, including swarming, making it an excellent model to investigate cooperative group behavior and its evolutionary stability [8]. Swarming is a collective movement performed by a variety of organisms from insects, fish, birds to microbes [9]. In bacteria swarming is a cooperative group motion by which cells migrate rapidly over surfaces, forming dynamic patterns of whirls and jets [10]. Cooperative movement is dependent on secreted surfactants, which are needed in addition to flagella, to propel bacterial groups efficiently over surfaces [11]. *B. subtilis* swarmer cells secrete a lipopeptide antibiotic surfactin, a powerful surfactant [12, 13], which reduces surface tension and enables cells to propel across surfaces [11]. Surfactin production is governed by the *srfA* operon [14, 15], which is regulated by quorum sensing [16–18]. Producers of surfactin release this public good into the surrounding environment, benefitting both the surfactin-producing cells and non-producing cells in the population[15, 19, 20]. However, the avoidance of metabolic investment by non-cooperative individuals can lead to their additional benefits, potentially destabilizing cooperation [1, 2]. Indeed, it was recently shown that Δ*srfA* mutants gain a reproductive fitness advantage in a mixed swarm by exploiting extracellular surfactin produced by the isogenic wild type strain and thus behave as cheats [19]. However, stability of public goods sharing during bacterial cooperative behaviors in general and cooperative swarming specifically, remains poorly understood. For example, surfactant sharing in the presence of cheats (Δ*srfA* mutants) during repeated growth cycles has not been addressed experimentally thus far.

*B. subtilis* isolates are known to engage in kin discrimination like behavior [21]. As we have previously shown swarms of closely related *B. subtilis* strains (kin, 99,93-99,99 % average nucleotide identity (ANI)) merge upon encounter, but less related (non-kin, 98,73-98,84 % ANI) strains form visible boundary lines between their swarms [21, 22] suggesting that *B. subtilis* swarmer cells preferentially cooperate with their clonemates and close kin but avoid non-kin. Moreover, kin strains always formed a common swarm, while non-kin excluded each other by the positive frequency dependent selection mechanism [19]. *B. subtilis* isolates also engaged in kin discrimination like behavior in floating biofilms whereby kin strains again remained closely intermixed, while non-kin segregated into well visible patches with one strain gaining dominance over the other [23]. These results support prediction that kin discrimination mechanisms may be important for evolutionary stability of cooperative behaviors. However, despite progress in our understanding of bacterial kin discrimination, several knowledge gaps persist in this area. One such gap pertains to the role of kin discrimination in limiting the spread of cheats in evolved populations.

To address these critical questions, we employed experimental evolution approaches. Our results demonstrate that the advantage observed in surfactin cheats during co-swarming with isogenic producers is not stable but rather quickly leads to swarming collapse due to the fitness advantage of cheats, resulting in a scenario akin to “the tragedy of the commons” [24, 25]. Furthermore, non-swarming strains exhibited lower competitive index when mixed swarm composed of surfactin cheats and isogenic surfactin producer was staged against antagonistic non-kin strain, than when staged against isogenic or kin strain, suggesting that kin discrimination associated interactions hinder the advantage of Δ*srfA* mutant strain. Finally, we show that the surfactin cheats evolve disproportionately more frequently during experimental evolution involving cooperative kin interactions, than during evolution involving antagonistic non-kin interactions, which indicates that differential behavior towards kin versus non-kin has consequences for social evolution and stability of cooperation. Moreover, our results highlight the potential role of kin discrimination as a mechanism acting against cheats spread within species.

## METHODS

### Strains and media

Strains used in this study are described in Table 1. Strain BM1658 was prepared by transforming strain BM1336 [21] with pMS17 plasmid DNA (EM1096) [26] to obtain a strain with constitutively expressed yellow and red fluorescent proteins and two antibiotic resistances.

**Table 1:** Strains and plasmids used in this study.

| Strain | Strain abbreviation | Genotype | Reference |
| --- | --- | --- | --- |
| BM1658 | PS-216 $\text{Kn}^R$ $\text{Sp}^R$,<br>focal strain | PS-216 *sacA::p43-mKate2* ($\text{Kn}^R$)<br>*amyE::p43-yfp* ($\text{Sp}^R$) | This work |
| BM1629 | PS-216 $\text{Kn}^R$ | PS-216 *sacA::p43-mKate2* ($\text{Kn}^R$) | Špacapan et al., 2020) |
| BM1345 | PS-216 $\text{Sp}^R$ | PS-216 *amyE::p43-cfp* ($\text{Sp}^R$) | Stefanic et al., 2021 |
| BM1347 | PS-218 $\text{Sp}^R$ | PS-218 *amyE::p43-cfp* ($\text{Sp}^R$) | Stefanic et al., 2021 |
| BM1328 | PS-216 $\text{Cm}^R$ | PS-216 *sacA::p43-yfp* ($\text{Cm}^R$) | Stefanic et al., 2021 |
| BM1044 | PS-216 $\Delta srfA$ | PS-216 *srfA::Tn917* ($\text{MLS}^R$) | Oslizlo et al., 2015 |
| PS-216 | PS-216 | Wild type | Stefanic & Mandic-Mulec, 2009 |
| PS-13 | PS-13 | Wild type | Stefanic & Mandic-Mulec, 2009 |
| PS-218 | PS-218 | Wild type | Stefanic & Mandic-Mulec, 2009 |
| BM1336 |  | PS-216 *amyE*::p43-yfp (Sp<sup>R</sup>) | Stefanic et al., 2015 |
| EM1096 | | *Escherichia coli* DH5 $\alpha$ (Amp<sup>R</sup>)<br>with plasmid pMS17<br>(*sacA*::P43-mKate2 (Kn<sup>R</sup>) ) | Špacapan et al., 2020 |

Strain abbreviation names (Table 1) indicate the name of the parental wild type strain and the specific antibiotic resistance and will be referred to in the text as wild type strains. Mutant strain BM1044 [18], which does not produce surfactin will be referred to as PS-216 Δ*srfA*.

Bacteria were cultivated in LB (Lennox) agar with 2 % agar and LB (Lennox) liquid media supplemented with antibiotics (Kanamycin 25 μg/ml – Kn, Spectinomycin 100 μg/ml – Sp, Chloramphenicol 10 μg/ml – Cm or MLS (Erythromycin 0,5 μg/ml and Lincomycin 12,5 μg/ml). Swarming agar also known as B media was used for swarming experiments and was prepared as previously described by Stefanic and Belcijan et al. (2021) with final agar concentration of 0,7 % [22].

M9 media was used to grow cultures of isolated evolved strains for whole genome sequencing. M9 media was prepared by sterilization of 1x M9 salts and addition of sterile solutions of MgSO_4_ x 7H_2_O (5 mM), CaCl_2_ (0,2 mM), glucose (1 %), trace metal mix (0,25 x trace metal mix: 12, 5 μM FeCl_3_, 5 μM CaCl_2_, 2,5 μM MnCl_2_ x 4H_2_O, 2,5 μM ZnSO_4_ x 7H_2_O, 0,5 μM CoCl_2_ x 6H_2_O, 0,5 μM CuCl_2_ x 2H_2_O, 0,5 μM NiCl_2_ x 6H_2_O, 0,5 μM Na_2_MnO_4_ x 2H_2_O, 0,5 μM Na_2_SeO_3_ x 5H_2_O, 0,5 μM H_3_BO_3_), BME vitamin mix (0,25 x), sodium glutamate (0,04 %) and casein hydrolysate (0,25 %).

### Swarming assay

Each swarming plate was prepared by pouring exactly 15 mL of swarming medium onto a Petri dish and allowed to cool upright overnight. Overnight cultures were prepared from frozen cultures, which were first plated, and single colonies used to prepare overnight culture. Overnight cultures were grown for 16 h at 37°C in LB or LB with an antibiotic. Plates inoculation was carried out by spotting a 2 μL drop of culture onto the swarming agar plate and allowed to dry. Plates were then incubated at 37°C and 80 % relative humidity (RH) for 22-24 h for swarms to form.

### Temporal dynamics of co-swarming surfactin producing and nonproducing strains

We determined the temporal dynamics of the relative frequency of non-swarming PS-216 Δ*srfA* (MLS^R^) mutant strain that does not produce surfactin after co-swarming with the wild type PS-216 strain on the swarming agar over three cycles of reinoculation. Overnight cultures of PS-216 and non-swarming mutant strain PS-216 Δ*srfA* (MLS^R^) were mixed 1:1. and their concentration in a mixture verified by CFU plating on LB agar with selection for antibiotic resistance. Swarming assay was performed and after each re-inoculation cycle, we sampled 20 agar samples at the swarm edge using trimmed pipet tips. Agar samples with cells were resuspended in 250 μl saline solution (0,9 % NaCl solution), vortexed vigorously and resuspended cells (2 μL) reinoculated onto the centre of the fresh swarming agar. The number of cells in a sample was determined by CFU counts on LB agar or LB agar supplemented with MLS. Cycle of sampling and reinoculation were repeated until no swarming was observed on the plates after 24 h. To determine the temporal dynamic of the ratio between non-swarming PS-216 Δ*srfA* (MLS^R^) mutant strain and swarming wild type PS-216 strain at the edge of a common swarm, the relative frequency of each strain was monitored in each cycle. The experiment was repeated in three independent experiments, each in six evolutionary trajectories. Heatmap was produced using OriginPro 2018 (Northampton, USA).

### Competitive index of the *srfA* mutant during co-swarming with its wild type strain

We determined the competitive index of the focal mutant PS-216 Δ*srfA* in a co-swarm with the wild type PS-216 Kn^R^ strain after the contact with either isogenic kin (PS-216 Sp^R^) or non-kin (PS-218 Sp^R^) strain. Competitive index is defined as output ratio between the two strains divided by their input ratio [28, 29] and is used to determine the growth of the mutant strain according to wild type strain during mixed colonization of prospective niches [30].

Overnight culture of PS-216 Δ*srfA* (MLS^R^) and PS-216 (Kn^R^) were mixed in 1:1 ratio and the relative frequency of each strain was determined by plating on LB agar with antibiotics according to resistance of each strain (MLS or Kn, respectively). CFUs were counted after overnight incubation at 37°C. During swarming assay, 2 μl of the mixed culture was inoculated opposite kin PS-216 Sp^R^ strain or non-kin PS-218 Sp^R^ strain (3 cm apart) (Figure 2A). We sampled 20 agar cores using trimmed 1 ml pipet tip at the area where mixed swarm met the opposing swarm (PS-216 Sp^R^ or PS-218 Sp^R^). The samples were resuspended into 250 μl of saline solution (0,9 % NaCl). Samples were vigorously mixed, and the relative frequency of co-swarming strains were again determined by plating on LB agar with selection for antibiotic resistance (MLS or Kn, respectively). We determined the competitive index (CI) of the focal strain PS-216 Δ*srfA* at the boundary with non-kin strain or the merging point of kin strains. The competitive index (CI) was calculated by dividing *R*_*f*_ with *R*_*i*_ as indicated in Equation 1. The initial ratio between the two co-swarming strains (PS-216 Δ*srfA* and PS-216 Kn^R^) (*R*_*i*_) was determined in the inoculum and the final ratio (*R*_*f*_) at the swarm meeting area after swarming cycle was completed (Equation 1).

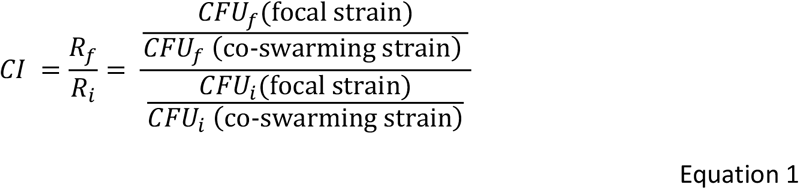

**Figure 1:**
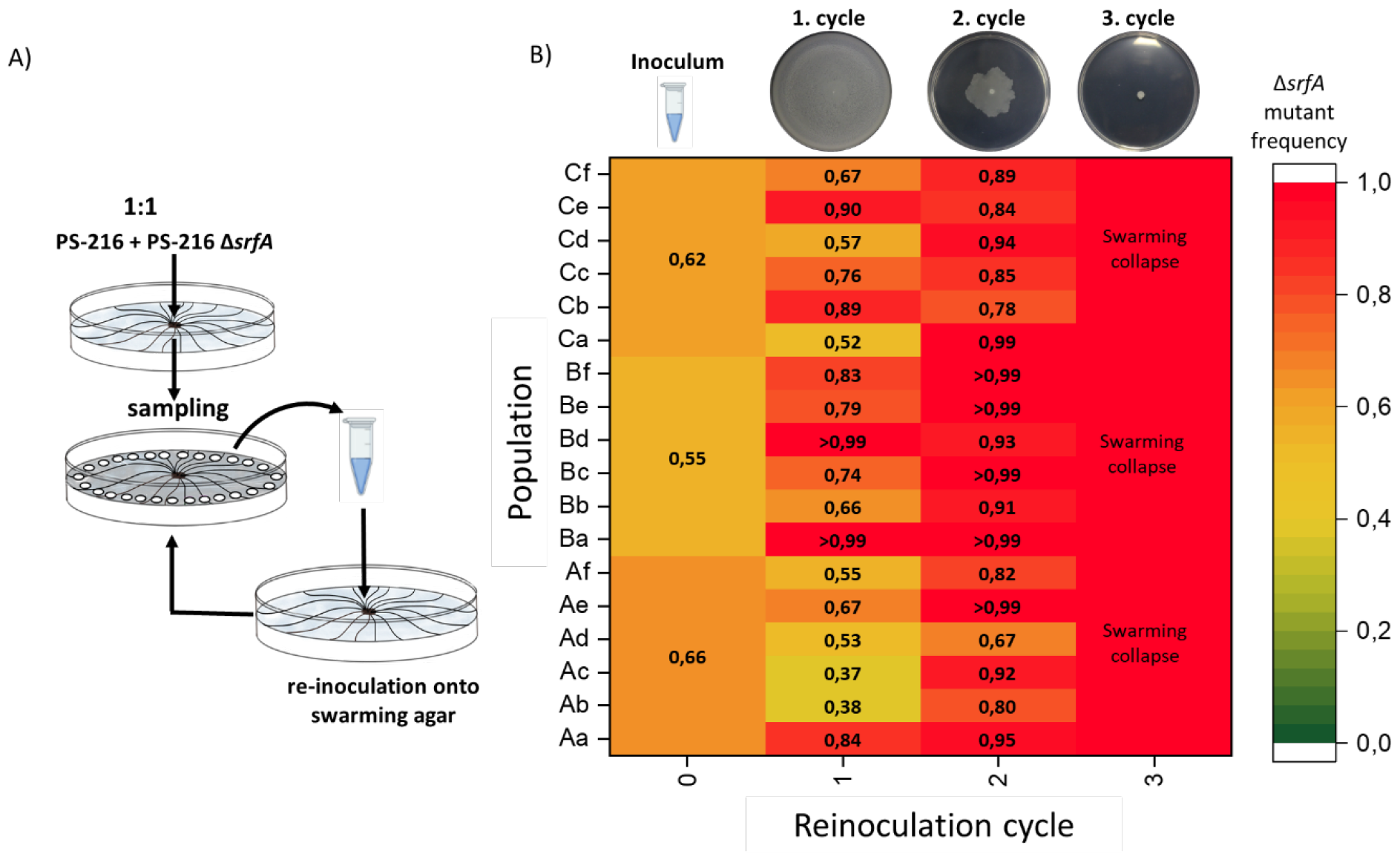
The dynamics of surfactin mutant PS-216 frequencies during co-swarming with the wild type strain PS-216 on swarming agar. A) Schematic representation of the experiment. B) The heatmap represents the PS-216 ΔsrfA strain relative frequency in the entire population. The first column represents the PS-216 ΔsrfA relative frequency in the inoculum and the successive columns the PS-216 ΔsrfA relative frequency in subsequent reinoculation cycles until population was not able to swarm (swarming collapse). We performed three independent experiments (A-C), each in six trajectories (a-f). Increasingly red squares represent the increasing relative frequency of surfactin mutant strain. The representative images of plates after each cycle of reinoculation of co-swarming strains are positioned above the heatmap.

**Figure 2:**
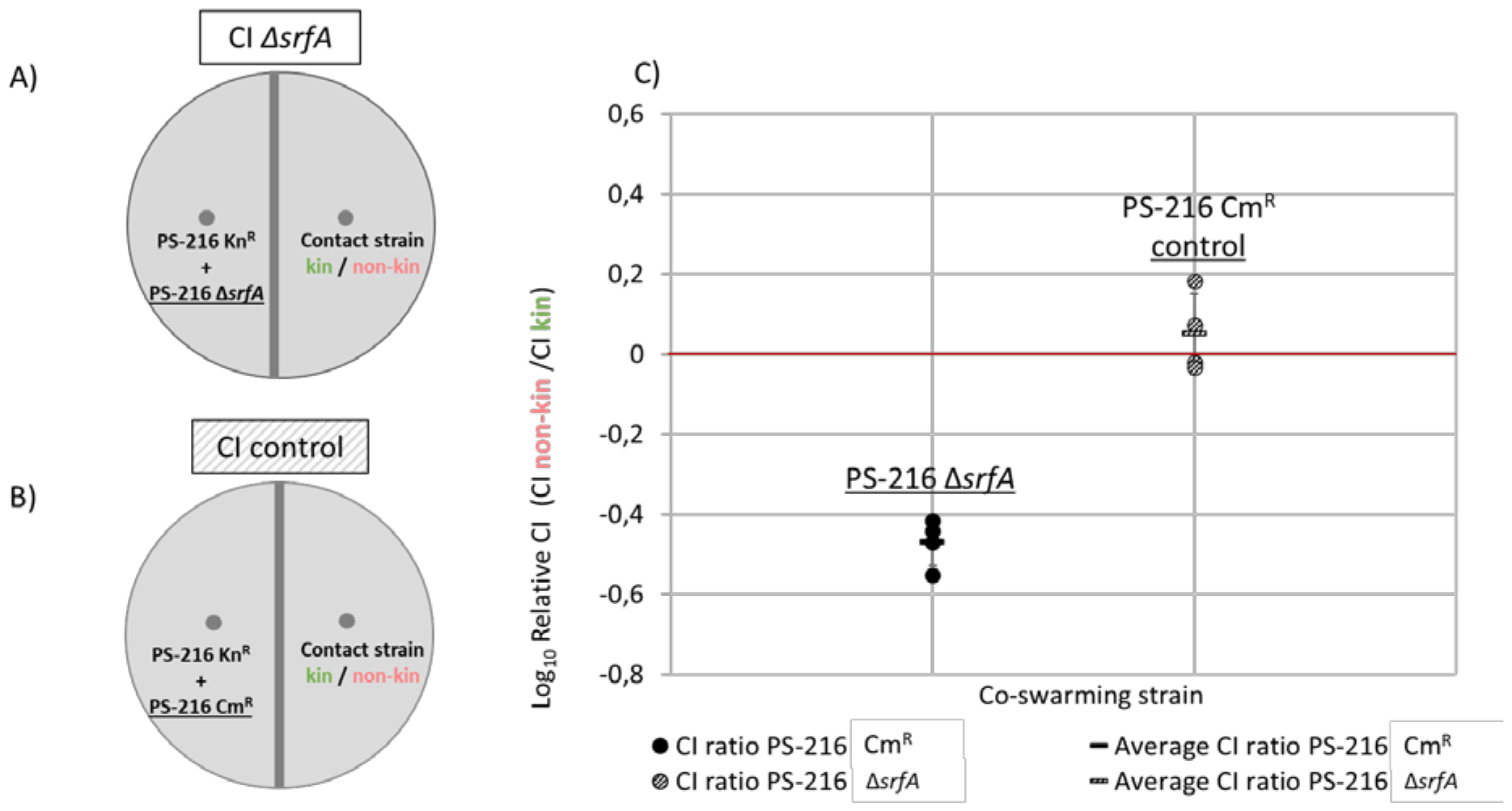
Relative competitive index of surfactin mutant or wild type strain in mixed swarm after contact with kin or non-kin swarm. A, B) Schematic representation of the experiment where A) surfactin non-producing mutant (PS-216 ΔsrfA) or B) swarming strain PS-216 Cm^R^ was mixed with isogenic wild type strain (PS-216 Kn^R^) in 1:1 ratio and inoculated on one side of the agar opposite to the isogenic kin strain (PS-216 Sp^R^) or non-kin (PS-218 Sp^R^) strain and allowed to swarm for 24 hr. We sampled at the swarm encounter area to determine CFU of each strain in the mixed swarm and calculated the competitive index (CI). C) Relative CI represents the ratio between CI of focal strain (either PS-216 or PS-216 ΔsrfA, underlined in schematic representation A and B) when mixed swarm was staged against non-kin strain (CI_PS-218_) and it’s CI when mixed swarm was staged against kin strain (CI_PS-216_) (Equation 2). The ratios were consistent throughout independent experiments. Relative CI was calculated for each of four independent experiments, each performed in three replicates. Error bars represent the standard deviation (SD) of four repeats. Difference between the calculated relative CI of the swarming mixture with the srfA mutant compared to the control experiment with two wild types is statistically significant (p < 0,05).

As a control experiment, we co-inoculated two swarming strains, focal strain PS-216 Cm^R^ and co-swarming strain PS-216 Kn^R^, and again staged the co-swarm against kin strain (PS-216 Sp^R^) or non-kin strain (PS-218 Sp^R^) on a swarming media (Figure 2B) followed by CFU determination as described above. As described above we determined the ratio between focal strain PS-216 Cm^R^ and co-swarming PS-216 Kn^R^ in the inoculum and at the meeting area with a contact strain (kin PS-216 strain or non-kin PS-218 strain). Competitive index (CI) of a PS-216 Cm^R^ strain was calculated (Equation 1). The cell numbers of contact strain cells were disregarded.

Due to high variability between CI for the focal strains (PS-216 Δ*srfA* or PS-216 Cm^R^) in four independent experiments (Supplementary Fig. 3), we determined the relative CI as the ratio between CI of the focal strain when mixed swarm was staged against non-kin strain (CI_PS-218_) and it’s CI when mixed swarm was staged against kin strain (CI_PS-216_) (Equation 2). The ratios were consistent throughout independent experiments.

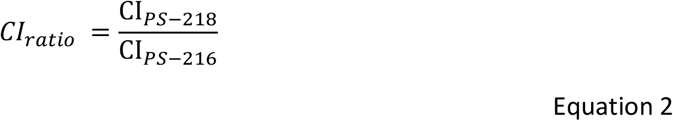

Both experiments were performed in four independent experiments, each in three replicates. T-test was used for statistical analysis and p value < 0,05 was presumed as statistically significant difference (Microsoft® Excel®).

### Experimental evolution of strain PS-216 in contact with isogenic, kin, or non-kin strain

Experimental evolution of the focal strain PS-216 Sp^R^ Kn^R^ was performed in 3 steps as presented in Figure 3A: A) Swarming assay was performed by inoculation of swarming agar (B media) with the focal and contact strain (isogenic PS-216, kin PS-13 or non-kin PS-218 strain) at opposite sides of the agar (∼ 3 cm apart) and both strains were allowed to swarm across agar overnight at 37°C and controlled humidity (80% RH). B) Next day sampling of bacterial cells at the swarm encounter area was performed, by coring 20 agar samples using trimmed pipet tips. The agar cores were resuspended in 0,5 ml of 0,9 % saline solution (0,9 % NaCl). C) The focal population’s cells were selected by growing (2 % inoculum) in liquid LB media supplemented with antibiotics spectinomycin (Sp) and kanamycin (Kn) for 3 h. After selection swarming assay was repeated by reinoculation of the focal populations (2 μl) on swarming agar opposite to a contact strain, as described under A. Populations sampled in the swarm encounter area were frozen in 7 % glycerol at -80°C for later analyses. In total 20 cycles of re-inoculation and exposure to contact strain (PS-216 (self), PS-13 (kin), PS-218 (non-kin)) were performed in 6 trajectories for each contact strain.

**Figure 3:**
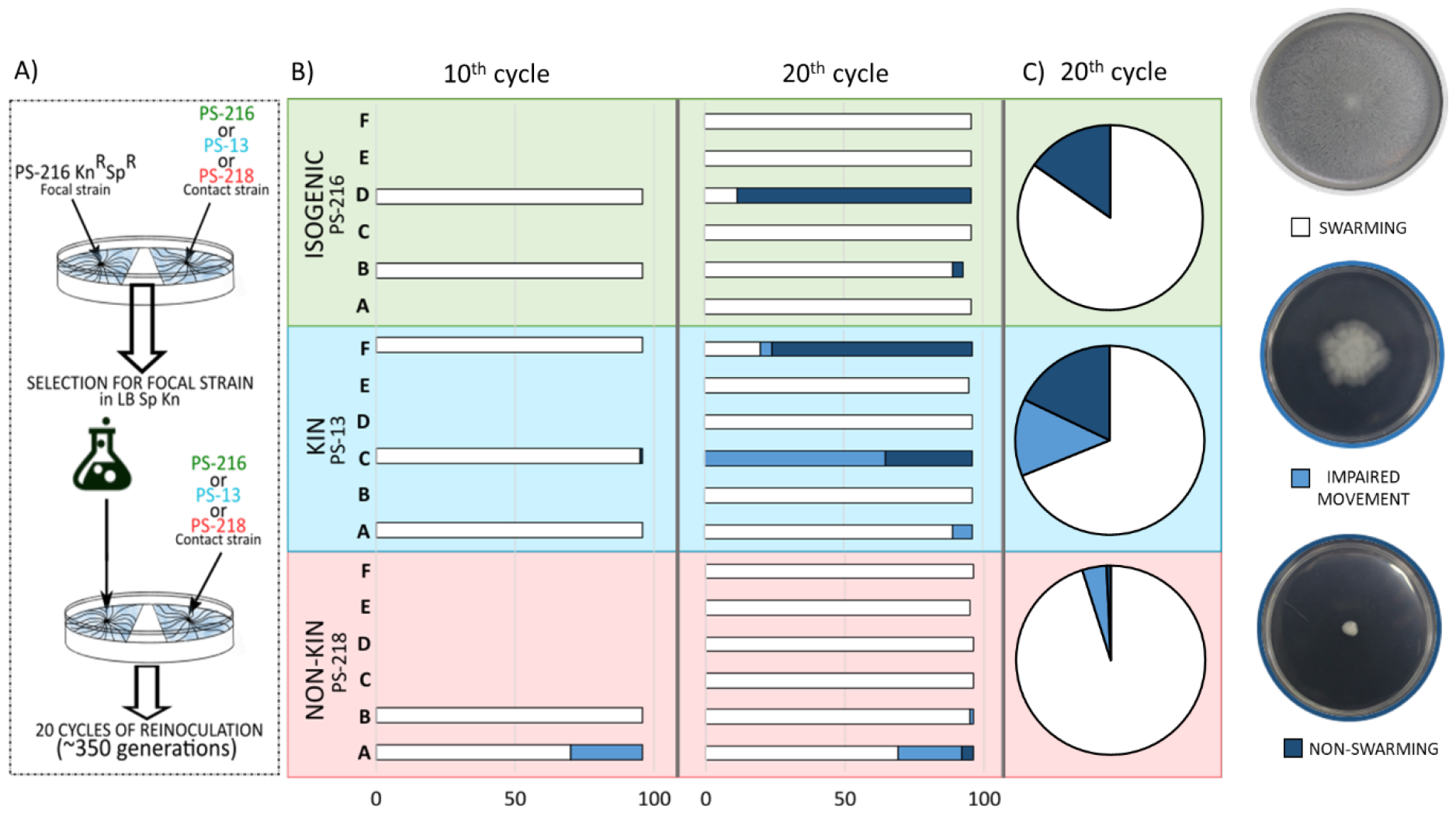
A) Scheme of experimental evolution of B. subtilis strain PS-216 Kn^R^ Sp^R^ which was periodically exposed to contact strain (isogenic PS-216, kin PS-13, non-kin PS-218) on swarming agar. At the contact point of two swarms the evolved population was sampled and selected in LB with antibiotics and used for the next inoculation. The cycle was repeated 20 times. B) Swarming phenotypes of randomly selected clones (∼ 96 clones per trajectory) were isolated from 7 evolved populations at the 10^th^ cycle and 18 evolved populations at 20^th^ cycle that were in periodic contact with isogenic strain (PS-216, green), kin strain (PS-13, blue) or non-kin strain (PS-218, pink). Swarming phenotype of each clone was determined, and columns are coloured accordingly for each evolved population – clones capable of swarming across entire swarming agar plate (70 mm in dimeter) in white; clones with impaired movement (small swarm without dendrites) in light blue; non-swarming clones in dark blue. Figures above the graph are exemplifying all three swarming phenotypes. C) Overall percentage of each phenotype found in all six trajectories exposed to the same contact strain.

### Screening for swarming defective phenotype in evolved populations

Evolved populations of the final (20^th^) evolutionary cycle (six populations for each of three contact strains PS-216, PS-13, and PS-218) were screened for swarming defective phenotype. Total of 18 frozen populations (six for each of three contact strains) were scraped into 100 μl of saline solution (0,9 % NaCl), diluted up to 100 times, spread on the LB agar media supplemented with kanamycin and spectinomycin (LB Sp Kn) and incubated over night at 37°C for single colonies to grow. Approximately 98 colonies were selected at random for each of 18 evolved populations and resuspended in 50 μl of liquid LB Sp Kn medium. 2 μl of this suspension was then inoculated onto the centre of swarming agar plates and incubated for 24 h at 37°C and 80 % RH to allow swarms to form. Evolved clones were defined as swarming if swarming phenotype was the same or very similar to the parental strain, as impaired movement if the clone was capable of movement but formed a smaller swarming colony without intricate dendritic pattern, and as non-swarming if a clone did not swarm on semisolid B medium and only formed a compact colony at the inoculation point.

### Drop-collapse test

The drop-collapse test was performed according to the protocols used by Bodour & Miller-Maier (1998) and Youssef et al. (2004). Briefly, frozen (−80 °C) evolved clones and parental cultures were revitalized on LB agar and then overnight cultures were grown from single colonies in liquid LB by shaking at 200 rpm at 37 °C for 16 h. Overnight culture (1mL) was centrifuged and spent media transferred into fresh centrifuge tubes and diluted with MQ water in 96-well plate. Methylene blue (2 μl) was added to 80 μL of each dilution to increase the visual contrast of droplets. 3-5 μl of mineral oil was pipetted onto the lid of 96-well microtiter plate and incubated et least 1 h at room temperature for oil to spread in each well. We pipetted 2 μl of spent media samples onto oil covered wells of the microtiter plate lid and after one-minute incubation formed droplets were imaged by using stereomicroscope (Leica WILD M10, Leica Mycrosystems, Inc., Danahe Corporation, Wetzlar, Germany). Droplet sizes were determined using ImageJ (ImageJ 1.53f51, Java 1.8.0_172 (64-bit), Wayne Rasband, National Institutes of Health, USA) with Vernier calliper used as scale for spatial calibration (determining pixel – distance ratio using Set scale function in ImageJ). Concentration of surfactants in spent media were determined according to standard curve obtained from measuring droplet sizes of known surfactin sodium salt (Fujifilm Wako Pure Chemical Corporation, Osaka, Japan) concentrations. Surfactin sodium salt was dissolved in PBS buffer and dilution (80 μg/mL to 0.04μg/mL) were prepared in MQ water. 2 μl of methylene blue was added to increase contrast of droplets. Droplet size was determined using ImageJ as described above and standard curve was drawn in Excel (Microsoft® Excel® for Microsoft 365 MSO (Version 2204 Build 16.0.15128.20158) 64-bit) and used to measure surfactant concentrations in spent medium of parental and evolved clones. Significant difference according to parental strain surfactant concentration was determined by the Man-Whitney U-Test for two independent samples (IBM SPSS Statistics, Version 29.0, IBM Corporation 1989, 2015, 64-bit).

### Sequencing

Seven evolved strains originating from evolved populations in contact with either isogenic, kin or non-kin were selected for sequencing. 21 evolved clones were revitalized from frozen cultures on LB agar plates and grown over night at 37°C. Single colony was inoculated into liquid LB media and grown at 37 °C for 16 h with shaking at 200 rpm. M9 medium was inoculated using overnight culture (1 % inoculum) and culture was grown to early exponential growth phase at 37 °C with shaking at 200 rpm (approximately 4 hours and OD_650 nm_ 0,2-0,3 a.u.). Obtained culture was pelleted by centrifugation (5 min, 10 000 g), media was removed, and cells washed with saline solution (0,9 % NaCl). Cells were pelleted once more by centrifugation (5 min, 10 000 g), supernatant was removed, and culture was frozen at -80 °C. Frozen culture samples were sent for DNA sequencing (Macrogen Korea, Geumcheon-gu, Seoul, Korea).

Because DNA of sufficient quality and quantity for whole genome sequencing (WGS) was not obtained from frozen culture samples of some evolved clones by Macrogene (Korea, Geumcheon-gu, Seoul, Korea), we isolated DNA from these strains using the following protocol. Single colonies of revitalized evolved strains were inoculated into liquid LB media and incubated for 16 h with shaking at 200 rpm at 37 °C. Culture was inoculated into fresh LB media supplemented with 1 % glucose and grown to early exponential phase (3,5 h at 37 °C with shaking at 200 rpm). DNA was isolated from 2 ml of bacterial culture using EURx Basic DNA Purification Kit (EURx Sp. Z o.o., Gdansk, Poland) following standard protocol. DNA was eluted twice using 50 μl of hot (80 °C) Elution Buffer. DNA quality control was performed using 4200 TapeStation System (Agilent, Santa Clara, CA, United States) by standard protocol using Genomic DNA ScreenTape and TapeStation Software (Agilent, Santa Clara, CA, United States) (kindly provided by prof. dr. P. Trontelj at the Department of Biology, Biotechnical Faculty, University of Ljubljana).

Sequencing of genomic DNA of 21 evolved clones and the parental strain was performed on Illumina HiSeq X10 platform (Macrogen Korea, Geumcheon-gu, Seoul, Korea) with libraries prepared with Illumina-Shotgun library (Illumina TruSeq DNA PCR-free (350 bp insert)), which generated 150 bp pair-end reads (genome sequences of evolved strains available in the NCBI database under BioProject accession number PRJNA1031373).

Parental strain PS-216 genome was previously sequenced using an Illumina MiSeq platform (genome sequence available in the NCBI database under BioSample accession number SAMN08637096) [22] and additionally sequenced by Pacific Biosciences RSII platform (PacBio) using 10 kbp single-molecule real-time (SMRTbell) library (Macrogen Korea, Geumcheon-gu, Seoul, Korea).

### Sequence assembly and analysis

Genome of the parental strain PS-216 was assembled using short Illumina sequences and long range PacBio sequences. Adapters of Illumina sequences, low-quality bases, and low-quality reads (Q<25, shorter than 70 bp) were removed using Trimmomatic version v0.39 [33]. FastQC version v0.11.9 [34] was used to confirm the quality of the trimmed reads. No reads were flagged as poor quality, no overrepresented sequences or Illumina adapters were present in Illumina sequences. Canu v2.2 [35] was used to remove low-quality bases in reads of long-range PacBio sequences of the parental strain PS-216. Assembly of parental strain PS-216 genome was performed *de novo* using Unicycler v0.4.9 [36] with default parameters and normal bridging mode. Quast 5.0.2 [37] was used to determine assembly quality and the number of contigs. The assembly of the parental genome resulted in one contig (4102284 bp). Annotation was performed by PGAP 6.4 (NCBI Prokaryotic Genome Annotation Pipeline) [38].

Quality short reads of evolved strains was first determined using FastQC version v0.11.9 [34] and Trimmomatic version v0.39 [33] was used to remove adapters, low quality bases and low-quality reads. Genome assembly of evolved strains was performed *de novo* using Unicycler v0.4.9 [36] with default parameters and normal bridging mode.

Changes in the sequences of evolved strains such as SNP’s, insertions, deletions and insertions or deletions of mobile genetic elements were identified using Breseq v036.1 [39], which maps the reads of the evolved strains onto the annotated reference genome (PS-216) using Bowtie 2 [40]. To distinguish between sequence gaps that are a result of genome changes during experimental evolution and sequence gaps caused by incomplete assembly due to sequence repeats in the genome, we compared two types of mapping to the reference genome PS-216. First, we mapped the evolved strains genomes onto the parental strain PS-216 genome using algorithm BWA-SW 0.7.17 [41], after which data was compared to mapping of short reads of evolved strains onto the reference genome obtained using Breseq. If short reads of evolved strains’ genomes mapped onto locations where gaps were discovered with BWA-SW, we presumed that the observed gaps are a consequence of imperfect assembly and not true gaps in evolved strain genome. Sequence analysis with Trimmomatic, FastQC, Unicycler and Quast were performed using Galaxy web platform, specifically public server at galaxytrakr.org [42], version 21.09. Analysis with PGAP, Breseq (Bowtie2), and BWA-SW were performed in Linux Ubuntu 20.04.6 LTS (Canonical Ltd., United Kingdom).

Statistical analysis of the mutation number was performed using Mann-Whitney nonparametric test to compare mutation number of 7 genomes according to each contact strain (isogenic, kin, non-kin) (IBM SPSS Statistics, Version 29.0, IBM Corporation 1989, 2015, 64-bit).

## RESULTS

### Collapse of swarming in kin populations occurs due to spread of surfactin cheats

The collapse of cooperative behavior can be attributed to the proliferation of cheats within a population. When surfactant producing strain is mixed with its surfactin-deficient variant (kin exploiter), the wild type strain compensates for the mutant’s inability to swarm effectively [19, 43]. Intrigued by the fact that the mutant displays heightened fitness during co-cultivation [19], we sought to investigate whether this exploitative dynamic would eventually undermine the cooperative swarming.

To probe into the temporal dynamics between co-swarming wild type and surfactin-deficient strain, we combined the PS-216 *ΔsrfA* mutant and surfactin-producing PS-216 wild type strain in equal proportions (1:1 ratio). This mixture was inoculated onto the center of the swarming agar. After 24h incubation samples were collected from the advancing swarm edge, and the composition ratio of each co-swarming strain was determined by selective plating. This sampling and re-inoculation cycle was iterated until the point of swarming collapse (Figure 1A).

Upon the initial swarming cycle, the resultant mixed swarms managed to establish themselves on the surface of the swarming agar, in accordance with earlier findings [19, 44]. However, a subtle blurring of dendritic structures was observed within these swarms. In the second cycle the mixed swarm exhibited diminished size and lacked the well-defined dendrites characteristic of the PS-216 monoculture swarm. Progressing to the third cycle, a dramatic collapse of swarming occurred, with cells becoming immobilized at the initial inoculation point (Figure 1B). Notably, the proportion of the Δ*srfA* mutant within the mixed swarm increased with each cycle dominating the populations after just the second cycle (averaging 90 % of the total population) (Figure 1B).

### Local non-kin interactions hinder the advantage of *srfA* mutant strain limiting its spread in cooperative groups

According to theory kin discrimination can restrain exploitation and foster cooperation [6, 7, 45]. As established in previous studies of *B. subtilis* this mechanism can operate through territorial exclusion and antagonism between non-kin [19, 20, 45]. Therefore, it could also have a role in prevention of surfactin exploitation. To test the hypothesis that non-kin strains can antagonize non-producing mutant and prevent public good exploitation we evaluated the competitive index of PS-216 Δ*srfA* surfactin exploiter in a mixed swarm (PS-216 Δ*srfA* + PS-216 Kn^R^) upon encounter with genetically homogeneous swarm of isogenic cells (PS-216 Sp^R^) or a swarm of non-kin cells (PS-218 Sp^R^) (Figure 2A). The initial step involved preparing the inoculum by mixing PS-216 Δ*srfA* mutant and the wild type strain PS-216 at 1:1 ratio. The relative frequency of each strain within the inoculum was determined by CFU method to establish their concentrations in the mixture. This mixture was inoculated on the swarming agar opposite the inoculum of the contact strain (kin PS-216 Sp^R^ or non-kin PS-218 Sp^R^) (Figure 2A). Next day, the samples were taken at the swarm encounter area where the relative frequency of both co-swarming strains was determined. To assess the reproductive advantage of the mutant strain in mixed colonization relative to the wild type strain, the competitive index (CI) of the PS-216 Δ*srfA* mutant was calculated by dividing the final ratio between the two co-swarming strains by their initial ratio (Equation 1, Figure 2C).

Analyzing the competitive index (CI) of the *ΔsrfA* mutant within the mixed swarm when exposed to either kin or non-kin swarm unveiled a notable difference. The CI for the PS-216 Δ*srfA* mutant within the mixed swarm exhibited a significantly lower value when exposed to non-kin swarm compared to kin encounter (Supplementary Fig. 3B, Figure 2C) (p < 0,05). In contrast, we observed no statistically significant difference in CI of the control surfactin producing strain PS-216 Cm^R^ in mixed swarm (PS-216 Cm^R^ + PS-216 Kn^R^) when subjected to either isogenic kin (PS-216 Sp^R^) or non-kin (PS-218 Sp^R^) encounter (Supplementary Fig. 3B, Figure 2C; relative CI of PS-216 Δ*srfA* – black dots and control PS-216 Cm^R^ – striped, black dots).

Despite fluctuations observed in computed CIs across separate experiments (Supplementary Fig. 3B), the relative CI values (calculated as CI in non-kin interactions relative to CI in kin interactions) (Equation 2) exhibited consistency across independent experiments (Figure 2C).

### Antagonistic interactions limit the rise of surfactin non-producers during experimental evolution

Previous results suggest that non-kin *B. subtilis* strains engage in antagonistic behaviors [19, 21–23]. As result the process of kin discrimination may bear negative consequences for the emergence or proliferation of cheats, potentially shaping their evolution and aiding in stabilization of cooperative swarming behavior. To test the hypothesis, we conducted an experiment where the focal PS-216 Kn^R^ Sp^R^ swarming population was periodically exposed to isogenic (PS-216), kin (PS-13) or non-kin (PS-218) swarms at the designated swarm encounter area (Figure 3A). We predicted that mutants defective in surfactin production and consequently impaired swarming capabilities would appear less frequently in evolutionary trajectories associated with non-kin encounters.

Experimental evolution of PS-216 Kn^R^ Sp^R^ was performed on swarming B medium in six replicate populations (trajectories) for each experimental variant (A-F) for 20 re-inoculation cycles - equivalent to approximately 350 generations. The number of generations during 24h growth in liquid LB medium averaged 19,3 ± 1,7 generations, while growth on B medium yielded an average of 17,5 ± 0,9 (Supplementary Fig. 1). Additionally, we determined the mutation rate (see Supplementary Methods 2 for detailed description) for the wild type strain PS-216 being 3,72 × 10^−9^ mutations per cell division in LB media and 4,28 × 10^−9^ mutations per cell division during growth on swarming media (Supplementary Fig. 2).

Non-swarming clones evolved in two out of six populations subjected to repeated encounters with isogenic (PS-216) or kin (PS-13) swarm, respectively. The overall percentage of non-swarming clones within isogenic and kin encounters was 17,9 % and 15,4 %, respectively (Figure 3C, dark blue). In contrast, the evolution of non-swarming clones was observed in only one out of six populations exposed to non-kin swarm (PS-218) with notably lower overall percentage of evolved non-swarming strains (0,7 %) (Figure 3C, dark blue).

Populations in contact with kin strain PS-13 evolved clones exhibiting an impaired swarming phenotype in three instances (13,2 % overall percentage of impaired swarming strains) (Figure 3C, light blue) with two of these populations also containing non-swarming clones. Populations interacting with non-kin PS-218 strain featured impaired swarmers in two evolved populations, but their frequency was low, accounting for 4,2 % of sampled evolved strains (Figure 3C, light blue). One of these populations also included non-swarming strains. Furthermore, the combined percentages of non-swarming strains and strains with impaired movement on swarming agar were again higher in populations in contact with isogenic or kin strain (17,9 % and 28,6 %, respectively) compared to populations in contact with non-kin strain (4,9 %).

Moreover, we examined the frequency of swarming pattern changes during the 10^th^ cycle of experimental evolution for populations that exhibited change in swarming phenotype by the 20^th^ cycle. Non-swarming strains were detected in low relative frequency in population C in contact with the kin strain PS-13 (Supplementary Fig. 4), and they had not yet evolved in other trajectories. Interestingly, clones with impaired swarming in the population A exposed to the non-kin strain PS-218 contact were relatively frequent at the 10th cycle of reinoculation (27 %), but their frequency didn’t increase by the 20th cycle of reinoculation (24 %) (Supplementary Fig. 4). These results are in accordance with our hypothesis, that non-kin interactions limit the rise of non-swarming or swarming impaired mutants (proliferation or spread of non-swarming genotype in the population). To further delineate genetic bases of the mutants, we proceeded with the whole genome sequencing and subsequent analysis.

### Mutations in the *srfA* operon are frequent in evolved populations with history of kin encounters

Defect in swarming can be a consequence of different deleterious mutations, including mutations affecting flagellar function, chemotaxis or surfactin production [46]. Among these only surfactin production is considered bona fide social trait, the other two being private traits associated with an individual cell [47, 48]. To identify the genetic causes of the non-swarming phenotype we performed whole genome sequencing of 21 randomly selected evolved strains. These strains exhibited either a change in swarming behavior or retained the parental swarming pattern. For each of the three types of evolved populations (isogenic, kin or non-kin) we selected seven clones for sequencing.

We found that two strains (clones - EV216-K54 and EV216-K105) that had lost the ability to swarm and were in contact with isogenic strain PS-216 had a deletion of 13 base pairs (bp) in *srfAB* (encoding surfactin synthase subunit 2 [49]). Furthermore, a mutation at position 2880 causing a stop codon in *srfAA* was identified in clone EV216-K6, which was capable of swarming but did not form dendrites typical for wild type PS-216 strain. Among strains isolated from populations interacting with the related strain PS-13, five carried a mutation in *srfAB* (EV13-K15, EV13-K27, EV13-K31, EV13-K57 and EV13-K41). Non-swarming clones EV13-K41 and EV13-57 carried single nucleotide polymorphism at position 1900 that caused stop codon in *srfAB* and strains EV13-K15, EV13-K27 and EV13-K31 had a missense substitution (Leu496Pro) in *srfAB*. Finally, strain EV218-K2 isolated from the population in contact with the non-kin strain (PS-218) also carried a deletion in *srfAB* (including *comS*) (Figure 4, Supplementary Table 1).

**Figure 4:**
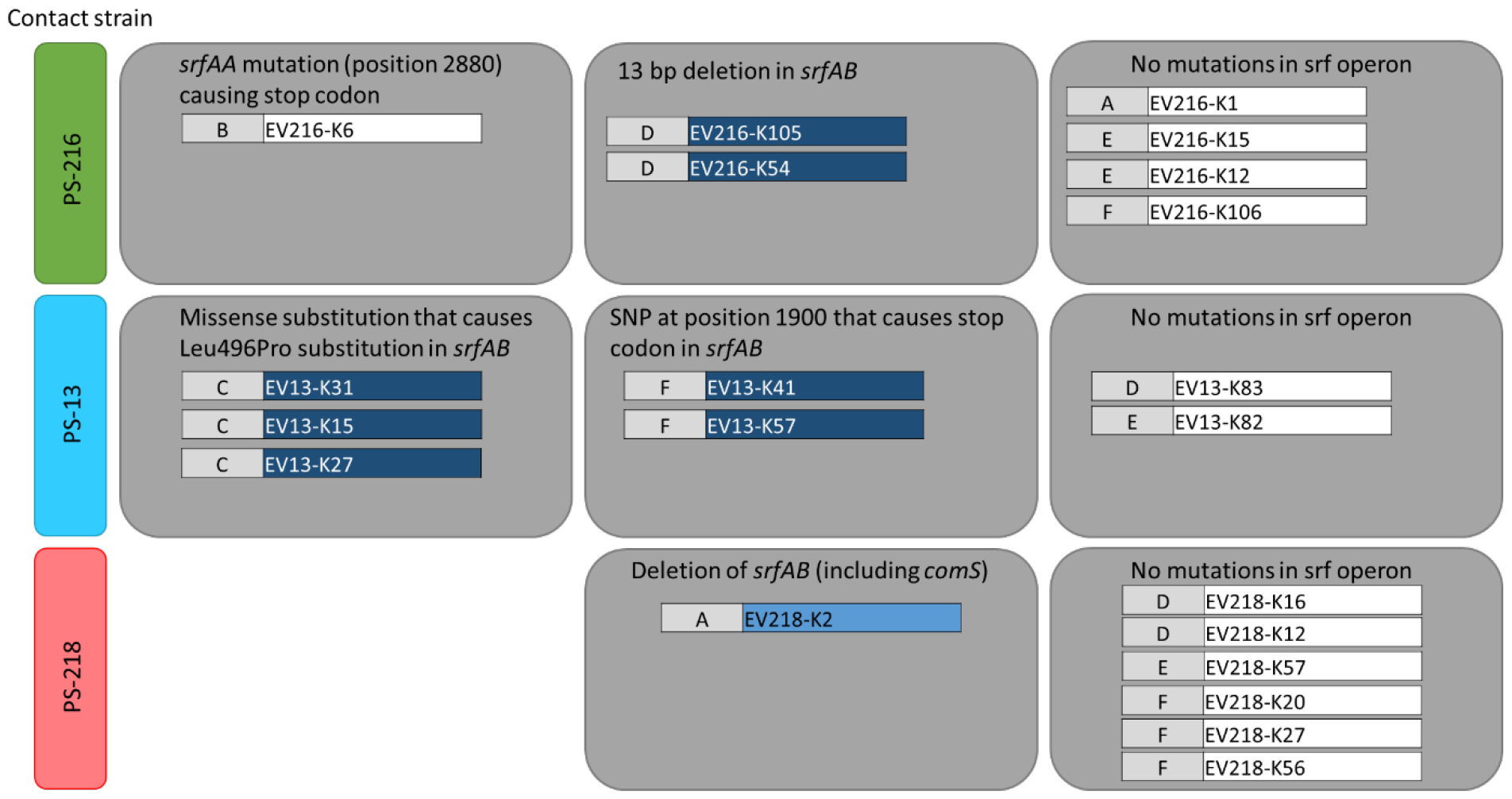
Sequenced clones and changes in genes important in swarming motility. Corresponding contact strain (isogenic strain PS-216, kin strain PS-13 or non-kin strain PS-218) and population from which they were isolated from (A-F) are stated on the left. The swarming phenotype of the evolved clone is represented by clone name background colour: white for swarming positive strains, light blue for strains with impaired movement on swarming agar that formed smaller swarms without distinct dendrites, and dark blue for clones unable to swarm on the surface of swarming agar. Gene mutations which might affect swarming motility are described in the upper part of each grey section.

Non-swarming strains within the same evolved population carried the same mutation, but those from different populations carried distinct mutations (Figure 4). This indicates that they evolved independently within each population, and we can also exclude cross contamination.

We did not detect a significant difference in overall number of mutations in sequenced strains regardless of the contact strain (isogenic, kin or non-kin) (p > 0,5). Altogether we found mutations in the *srfA* operon in 9 out of 21 sequenced genomes, among which only one originated from population with non-kin encounters. Although many genes contribute to swarming [46], none of the mutations outside the *srfA* operon (such as mutation in *epsE* involved in production of major biofilm matrix polysaccharide or response regulator *degU*) caused non-swarming or impaired swarming phenotype (Supplementary Table 1) [50, 51]. The average number of non-synonymous mutations per clone was 5 and did not significantly vary between clones from experimental evolution trajectories (Supplementary Table 1).

### Evolved clones with defective swarm phenotype show impairment in surfactant production

Sequencing of evolved clones revealed mutations in the *srfA* operon, which implied that non-swarming phenotype is caused by decreased surfactin production. Thereby we predicted that additional surfactin impaired mutants reside within each evolved population. To further verify this prediction, we randomly selected 20-24 clones from each of the three variants of experimental evolution (20^th^ reinoculation cycle) and tested them for the production of surfactin by a drop collapse test according to surfactin standard.

The number of evolved clones that produced less surfactin than the parental strain was again higher among strains isolated from the evolved populations in contact with isogenic strain (10 out of 24) and kin strain PS-13 (14 out of 21). In contrast, most evolved clones (18 out of 20) from populations in contact with non-kin PS-218 strain produced comparable quantities of surfactants as the parental strain PS-216 (p > 0,05) (Figure 5).

**Figure 5:**
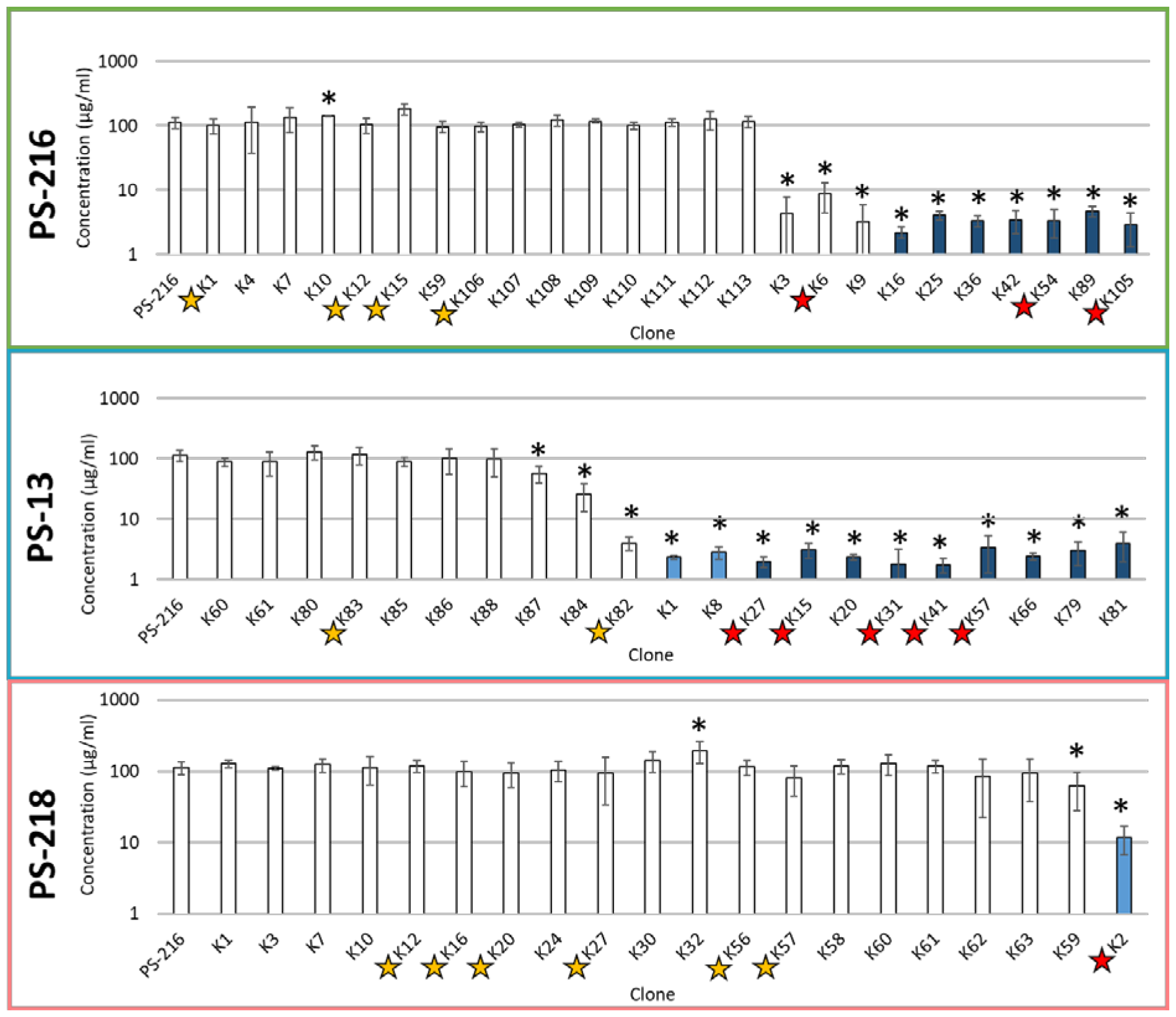
Surfactant concentrations of selected evolved clones in contact with isogenic strain (PS-216, green frame), kin strain (PS-13, blue frame) and non-kin strain (PS-218, pink frame). Graphs display produced surfactant concentrations of evolved clones (clone names are shortened according to Supplementary Table 2) in periodic contact with isogenic strain (PS-216), kin strain (PS-13) and non-kin strain (PS-218), respectively. Columns are coloured in white if evolved clones are able to swarm over entire swarming agar surface, in lighter blue if their movement is impaired (small swarm without dendrites) and non-swarming evolved clones in dark blue. Sequenced clones are marked with yellow star next to the clone’s name and the star is coloured red if clone carried a mutation in srf operon. Mann-Whitney U-test for two independent samples was performed and the clones that produced significantly different concentration of surfactin than the parental strain-wild type PS-216 (p < 0,05) are marked with the star (*) above the column.

All tested non-swarming clones (dark blue columns) produced significantly less surfactants than parental strain PS-216 (p < 0,05) (Figure 5). Most of the evolved clones without the change in swarming proficiency (white columns) produced similar levels of surfactants as the parental strain PS-216 (Figure 5). But there were few exceptions, such as EV216-K3, EV216-K6, EV216-K9, which evolved in contact with isogenic strain PS-216, and EV13-K82, EV13-K84, EV13-K87, which evolved in contact with kin PS-13 strain (white columns). These clones produced less surfactants but were still capable of swarming (Figure 5). Among these six indicated clones, the three clones (EV216-K3, EV216-K6, EV216-K9) colonized the swarming agar completely but did not form intricate dendrites pattern as the parental strain PS-216. One of the three evolved clones (EV216-K6) was also sequenced and found to carry the mutation in *srfAA*, which presumably lowered levels surfactin production, but still allowed swarming.

Overall, experimental evolution supports our hypothesis that non-kin interactions hinder the spread of mutants defective in surfactin production, which must rely on cooperative individuals to share this public good.

## DISCUSSION

Cheats exploit the work of others by consuming shareable resources that are costly to produce but benefit the entire population [2]. It is intriguing to understand what makes bacterial cooperation persist despite the presence of cheats. Here we experimentally demonstrate that kinship dependent interactions have an important role in maintaining cooperative behaviors. We show that when surrounded by kin cooperators the cheats, that do not contribute surfactin, quickly spread causing collapse of cooperation. However, the outcome changes if the cheats in the population are exposed to transient non-kin encounters, which significantly diminish surfactin mutant’s competitive index and hinder the proliferation of cheats in evolved populations.

Swarming motility of *B. subtilis* strains is supported by surfactin production, which acts as a public good [15, 43, 48]. Since public goods are shared and freely available, they can be easily exploited by cheats that benefit from public goods, but do not invest in their production [2]. Therefore, non-producers can outgrow producers in co-culture and cause “tragedy of the commons” [24, 25]. We show that surfactin cheats when mixed with the isogenic kin *B. subtilis* surfactin producers at 1:1 ratio quickly outgrows the producers and cause collapse of the collective swarming already at the 3rd inoculation cycle of the experiment. Previous work has revealed that surfactin non-producing strain can exploit surfactin produced by the wild type strain during the first swarm cycle [19, 44]. Non-swarming surfactin mutant strain also exhibited fitness advantage during co-swarming with the wild type strain on swarming agar [19, 44] and during sliding motility without flagella [48]. Our results are consistent with these findings, but also show that surfactant cheats can cause swarming collapse of swarming populations over rather short evolutionary time. This is in contrast to *Pseudomonas aeruginosa* swarming system, where the rise of surfactant cheats is restricted by metabolic prudence whereby the rhamnolipid production is limited only to stationary phase, where gain of fitness advantage of rhamnolipid cheater is not possible [52]. Moreover, *B. subtilis* NCBI 3610, which produces less surfactin than PS-216, is also better equipped to resist the spread of surfactin exploiters [44].

In contrast to cheater induced cooperation collapse of *B. subtilis* swarming, transient non-kin interactions restricted the fitness advantage of surfactin mutant co-swarming with the wild type surfactin producer. Additionally, non-kin interactions between two swarms limited the spread of *B. subtilis* swarming defective mutants that have arisen during experimental evolution. Specifically, non-swarming clones spread to higher frequency in evolved populations in contact with kin or isogenic swarm (15,4 % and 17,9 % clones with swarming ability loss, respectively) compared to evolved populations in contact with non-kin swarm, where only 0,7 % swarm defective clones were detected after 20^th^ cycle. Interestingly, in the earlier 10^th^ cycle, we also detected non-swarming strains only in one evolving population in contact with kin PS-13 strain where they were at low frequency and spread by the 20^th^ cycle. In contrast, one population exposed to non-kin strain PS-218 produced impaired spreaders already in 10^th^ cycle, nevertheless impaired swarmers failed to spread further by the 20^th^ cycle where their percentage was similar as in 10^th^ cycle. This was in striking contrast to the expansion of non-swarming strains in evolved populations in contact with isogenic (PS-216) or kin (PS-13) swarms. Based on these data we conclude that antagonistic non-kin interactions keep the surfactin exploiters at bay.

Sequencing of evolved non-swarming clones confirmed our prediction as it uncovered five distinct mutations in the *srfA* operon, specifically in genes *srfAA* and *srfAB*. As surfactin is needed for colonization of synthetic B medium (swarming agar) [14, 43, 53], swarming motility of such strains was impaired or even non-existent. Additionally, we detected significantly lower concentrations of surfactants in spent media of non-swarming evolved clones, suggesting that the detected mutations in the *srfA* operon are indeed responsible for non-swarming phenotype. These clones also appeared more frequently in self/kin exposed populations than in non-kin exposed populations.

The mechanism by which kin discrimination could limit the rise of non-swarming mutants remains unclear at this point. Lower competitive index of surfactin non-producing mutants and their failure to spread during experimental evolution could be brought about by different molecular mechanisms.

One of the explanations for limited spread of surfactin mutants can be an increased sensitivity of surfactin mutants to non-kin attack. Previous research suggested that surfactin producing strain accumulates stress phospholipid cardiolipin [54–56], which contributes to surfactin tolerance of *B. subtilis* [54]. The cytoplasmic membrane of the mutant *B. subtilis* lacking cardiolipin was observed to be more susceptible to the action of surfactin than the wild type [54]. Additionally, surfactin non-producing strain 168 halted its growth when inoculated onto surfactin containing agar compared to control without the surfactin[55]. However, the presence of surfactin in the growth medium of surfactin non-producing strain 168 also causes the changes in ratio between other phospholipid classes of the membrane that also increase resistance against surfactin induced permeabilization [55, 56]. Similarly, higher sensitivity to stress (osmotic, thermal and heavy metal stress) was also observed in *Pseudomonas aeruginosa* lacking exoproteases. This quorum sensing deficient cheats were also less resistant to oxidative stress, therefore proportion of cheats in the population, which is otherwise vulnerable to their invasion, decreased [57].

We have recently shown that kin discrimination promotes horizontal gene transfer (HGT) in *B. subtilis* through upregulation of competence genes important for uptake of extracellular DNA and transformation [22]. In addition, it was recently proposed by Lee et al. (2022), that horizontal gene transfer (HGT) is an important cooperation enforcement mechanism in bacterial populations. As a mutant cheater can arise in a cooperating population through a random loss-of-function, HGT can convert the cheater into a co-operator by reintroducing a lost cooperative gene and rescuing a cooperation [58]. Furthermore, it was also previously shown that HGT can act as a DNA repair mechanism [59, 60]. Therefore, elevated HGT between non-kin strains at the boundary between swarms, in relation to lower HGT between kin swarms, could contribute to the lower count of surfactin mutants at the boundary.

Differentiating kin from non-kin could also empower cooperators to sanction cheats in the population. Policing, mechanism that can enable cooperators to directly sanction cheats and thereby enforce cooperation [61], has been implicated for *P. aeruginosa*, which produces sharable protease along with a toxin called cyanide. As cooperators also generate an immunity factor safeguarding them against the harmful effects of cyanide, they are capable of effectively eliminating cheats. However, *P. aeruginosa* policing is directed towards cheats that arise within the same population. In contrast, *B. subtilis* policing might be triggered or even performed by phylogenetically non-kin conspecifics [62].

Here we provide strong support to the theory stating that the evolution of cheats is hindered by kin discrimination. Our research demonstrates that during evolution of *B. subtilis* on swarming agar random mutations in the *srfA* operon occur. Such mutants exhibit higher reproductive advantage than wild type and therefore swiftly spread in the population of surfactin producers. However, transient interactions with non-kin at the swarm encounter area hinder the advantage of surfactin non-producing mutants and restrict their proliferation which inadvertently benefits rival non-kin population. Therefore, described “reciprocity” between seemingly antagonistic populations might be involved in stabilisation of cooperation in microbial populations found in natural environment. Most importantly, we show that kin discrimination affects evolutionary outcome of cooperative behavior where non-kin, but not kin, swarms obstruct the rise of randomly emerging surfactin non-producing cheats during experimental evolution on swarming media.

## ACKNOWLEDGMENTS

This work has been financially supported by the Slovenian Research Agency (ARRS) program grant P4-0116 supporting the lab of IMM and by project grants J1-4411 (awarded to PS), J4-9302 and J4-4550 (awarded to IMM). We also acknowledge the infrastructural centre “Microscopy of biological samples” (Biotechnical Faculty, University of Ljubljana, Slovenia) for the use of stereomicroscope and prof. dr. P. Trontelj and his group (Department of Biology, Biotechnical Faculty, University of Ljubljana) for the use and help with TapeStation System. We would like to thank master students Zala Vašl and Jernej Kralj for technical assistance.

## Notes

### Competing Interest Statement

The authors have declared no competing interest.

