## Supplementary methods and figures for "The rise of cheats during experimental evolution is restricted by non-kin interactions between *Bacillus subtilis* soil isolates"

**SUPPLEMENT**

**Supplemental Methods**

Supplementary Method 1: Growth rate (the number of generations)

We determined the number of generations of the focal strain PS-216 that develop in liquid LB media or during growth on swarming media for 24 h. The strain PS-216 was revitalized from frozen (-80°C) culture and streaked onto LB agar plates and grown over night at 37°C. One colony was transferred into liquid LB medium and shaken (200 rpm) for 16 h at 37°C. Next day the culture was transferred into fresh liquid LB medium (1 % inoculum) and shaken (200 rpm) at 37°C for 3 h to reach exponential phase of growth. 2 µl of bacterial culture was inoculated in the center of the swarming agar and 3 µl of culture into 3 ml of liquid LB media. The number of cells in the inoculum was determined by spread plate method (CFUs per milliliter). Inoculated swarming agar plates were incubated for 24 h at 37°C with 80 % relative humidity (RH) and inoculated liquid LB media was shaken (200 rpm) for 24 h at 37°C. After incubation, the 1 ml of culture was sampled and cells on swarming agar were scraped from the agar surface into 2 ml of saline solution (0,9 % NaCl). We determined the number of cells by spread plate method (CFUs per milliliter) in the inoculum ($n_{inoculum}$) and after 24 h growth in liquid LB media or on swarming media ($n_{final}$). The number of generations was determined by using Supplementary Equation 1.

$$No. of\ generations = \frac{\log(n_{final}) - \log(n_{inoculum})}{log2}$$

Supplementary Equation 1

The experiment was carried out in three biological repeats, each performed in six replicates. The mean of three biological repeats and standard deviation were calculated.

Supplementary Figure 1

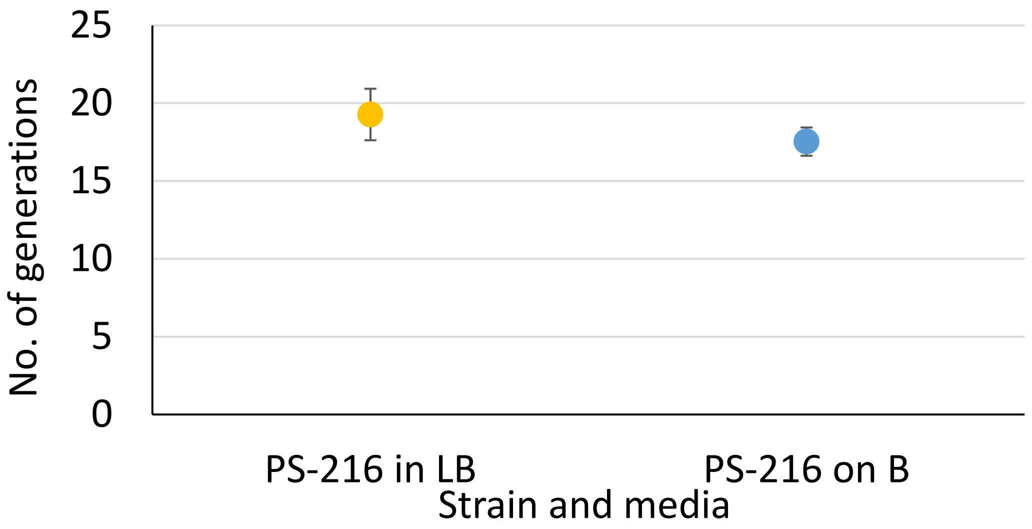


*Supplementary Figure 1: Number of generations of the focal strain PS-216 that grow in 24 h of growth in liquid LB medium or while swarming on swarming agar. Figure presents the mean with standard deviations of three independent experiments, each performed in six replicates as indicated in Supplementary Method 1. The error bars represent standard deviation of mean values of independent experiments.*

Supplementary Metnod 2: Mutation rate

We determined the mutation rate of focal strain PS-216 in liquid LB media and on swarming agar. The frozen culture (-80°C) of the focal strain was scraped and transferred into saline solution (0,9 % NaCl). The concentration of cells in the prepared cell suspension was determined by spread plate method (CFUs per milliliter). Cell suspension was inoculated at the center of swarming agar plates (2 µl) and into the liquid LB medium (3 µl). Inoculated swarming agar plates were incubated at 37°C with 80 % RH for 24 h, and inoculated liquid medium was shaken (200 rpm) at 24 h at 37°C. After overnight incubation we sampled 1 ml liquid culture and cells from the swarming agar were scraped and resuspended into 2 ml saline solution (0,9 % NaCl). The total number of cells in liquid media and on the surface of swarming agar after 24 h incubation was determined by spread plate method (CFUs per milliliter). By selection for antibiotic resistance against rifampicin ($Rif^R$) on LB agar media supplemented with rifampicin (5 µg/ml) we determined the total number of polymerase β subunit (*rpoB*) gene mutants which became resistant to rifampicin [1]. The experiment was carried out in three independent experiments, which were performed in six replicates. The mutation frequency was determined using web tool for Luria-Delbrük experiment - webSalvador [2]. We used Lea-Coulson Model with plating efficiency ε = 0,8, estimated final number of all cells (Nt) as the mean of total number of cells after 24 h of six replicates, initial number of mutations (Initial m) was set at zero and mutant counts were inserted. Using integrated algorithm, the point estimate number of mutations (m) in 1 ml culture, and the mutation rate (µ) were determined, including upper and lower limit of both variables.

Supplementary Figure 2

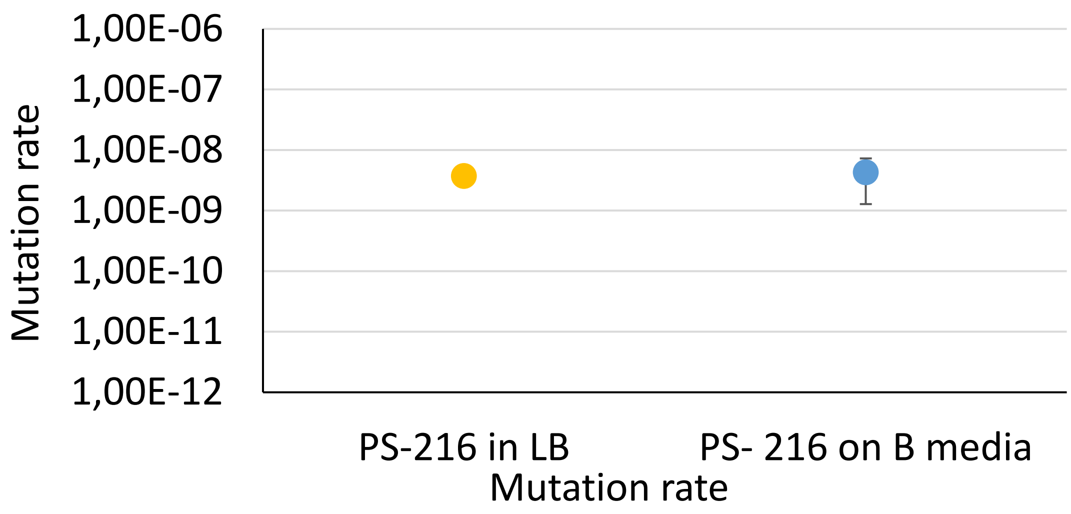


*Supplementary Figure 2: Mutation rate per cell per division determined by growing focal strain PS-216 in liquid LB medium or on semisolid swarming medium. Figure presents the mean with standard deviations of three independent experiments, each performed in six replicates as described in Supplementary Method 2.*

Supplementary Method 3: Competitive index of PS-216 Δ*srfA* mutant depending on kin or non-kin contact strain

We determined the competitive index of the focal mutant PS-216 Δ*srfA* in a co-swarm with the wild type PS-216 $Kn^R$ strain after the contact with either isogenic kin (PS-216 $Sp^R$) or non-kin (PS-218 $Sp^R$) strain.

Overnight culture of PS-216 Δ*srfA* ($MLS^R$) and PS-216 ($Kn^R$) were mixed in 1:1 ratio and the relative frequency of each strain was determined by plating on LB agar with antibiotics according to resistance of each strain (MLS or Kn, respectively). CFUs were counted after overnight incubation at 37°C. During swarming assay, 2 µl of the mixed culture was inoculated opposite kin PS-216 $Sp^R$ strain or non-kin PS-218 $Sp^R$ strain (3 cm apart) (Figure 2A) and allowed to swarm at 37°C with increased humidity (80 % RH) for 22-24 h. We sampled 20 agar cores using trimmed 1 ml pipet tip at the area where mixed swarm met the opposing swarm (PS-216 $Sp^R$ or PS-218 $Sp^R$). The samples were resuspended into 250 µl of saline solution (0,9 % NaCl). Samples were vigorously mixed, and the relative frequency of co-swarming strains were again determined by plating on LB agar with selection for antibiotic resistance (MLS or Kn, respectively). We determined the competitive index (CI) of the focal strain PS-216 Δ*srfA* at the boundary with non-kin strain or the merging point of kin strains. The competitive index (CI) was calculated by dividing $R_f$ with $R_i$ as indicated in Equation 1. The initial ratio between the two co-swarming strains (PS-216 Δ*srfA* and PS-216 $Kn^R$ ) ($R_i$) was determined in the inoculum and the final ratio ($R_f$) at the swarm meeting area after swarming cycle was completed (Equation 1).

As a control experiment, we co-inoculated two swarming strains, focal strain PS-216 $Cm^R$ and co-swarming strain PS-216 $Kn^R$, and again staged the co-swarm against kin strain (PS-216 $Sp^R$) or non-kin strain (PS-218 $Sp^R$) on a swarming media (Supplementary Fig. 2B) followed by CFU determination as described above. As described above we determined the ratio between focal strain PS-216 $Cm^R$ and co-swarming PS-216 $Kn^R$ in the inoculum and at the meeting area with a contact strain (kin PS-216 strain or non-kin PS-218 strain). Competitive index (CI) of a PS-216 $Cm^R$ strain was calculated (Equation 1). The cell numbers of contact strain cells were disregarded.

Both experiments were performed in four independent experiments, each in three replicates. T-test was used for statistical analysis and p value < 0,05 was presumed as statistically significant difference (Microsoft® Excel®).

Supplementary Figure 3

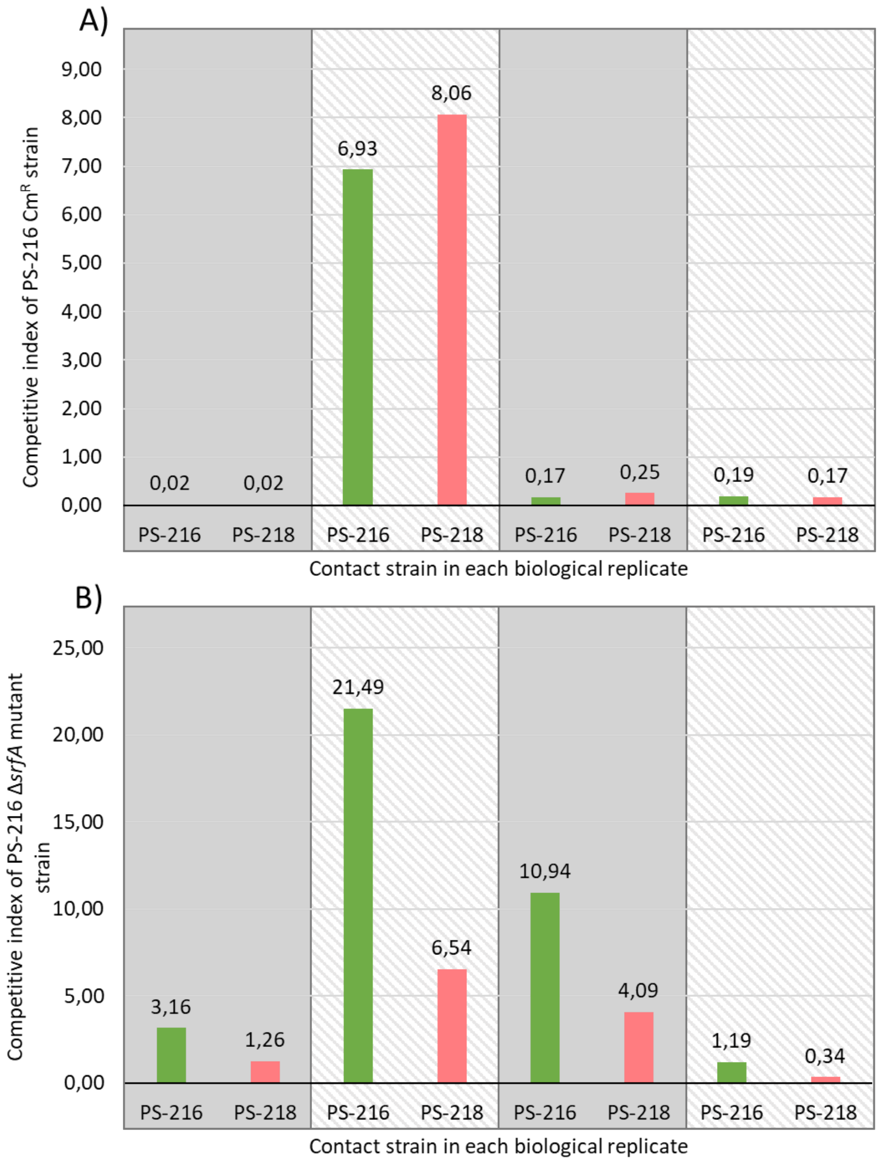


*Supplementary Figure 3: Competitive index (CI) of control strain PS-216 Cm$^R$ (A) or surfactin nonproducing mutant PS-216 Δ*srfA *(B) with all four individual experiments shown, each performed in three replicates as described in Supplementary Methods 3. (A) Competitive index (CI) of control strain PS-216 Cm$^R$ swarming in a co-swarm with w.t. strain PS-216 Kn$^R$ in contact with kin swarm PS-216 (green columns) or with non-kin strain PS-218 (pink columns). (B) Competitive index (CI) of non-swarming and surfactin nonproducing mutant PS-216 Δ*srfA *swarming in a co-swarm with w.t. strain PS-216 Kn$^R$ when in contact with kin strain PS-216 (green columns) or with non-kin strain PS-218 (pink columns).*

Competitive index (CI) of non-swarming and surfactin nonproducing mutant PS-216 Δ*srfA* swarming in a co-swarm with isogenic w.t. strain PS-216 Kn$^R$ was higher in all four biological replicates when common swarm was in contact with kin strain PS-216, than when a common swarm was in contact with non-kin swarm PS-218 (Supplementary Fig. 3B). Competitive index (CI) of control strain PS-216 Cm$^R$ swarming in a co-swarm with the isogenic strain PS-216 Kn$^R$ was similar regardless of the contact strain in all four biological repeats (Supplementary Fig. 3A)

<u>Supplementary Table 1: Table of all observed mutations in sequenced clones</u>

*Supplement table 1: Table of all evolved clones that were sequenced with observed mutations. Evolved clone's contact strain is indicated on the left and their swarming phenotype as a clone's background colour (swarming as white, impaired swarming in light blue and non-swarming in dark blue). Total number of mutations and all observed mutations are stated on the right side for each clone. Mutations in srfA operon are highlighted with green colour and mutations in eps operon are highlighted with yellow.*

| Contact strain | Strain | Population | Mutations | | | | | | |
|---|---|---|---|---|---|---|---|---|---|
| **PS-216** | **EV216-K1** | A | bmr3 (M351K) | oppA (Y142*) | yknY (Δ 1bp) | spoIIM (Δ 1bp) | | | |
| | **EV216-K6** | B | ywkF (G41A) | ywzG (Δ 7bp) | yxnA (G214A) | srfAA (W2280*) | tcyB (I40T) | queA (M130I) | Δ9 622bp (sipT, ykoA, ykpA, ykpB, ampS, ykpC, mreBH, abh, kinC) |
| | **EV216-K54** | D | mnmE (G327R) | srfAB (Δ 13bp) | yhfT (I380M) | pksL (A996E) | gltD (G194S) | yqjN (V265I) | levB (A282S) |
| | **EV216-K15** | E | yxcE (H135R) | ahpC (W170R) | resC (A64V) | | | | |
| | **EV216-K105** | D | mnmE (G327R) | srfAB (Δ 13bp) | yhfT (I380M) | pksL (A996E) | gltD (G194S) | yqjN (V265I) | levB (A282S) |
| | **EV216-K106** | F | glpK (A308V) | ywnA (C->T, -34) | | | | | |
| | **EV216-K12** | E | citH (G300D) | dctP (A367V) | slp (Δ 1bp) | yrkQ (F348Y) | licR (N203T) | yxkI (V147F) | |
| **PS-13** | **EV13-K15** | C | catD (A110V) | srfAB(L496P) | katA-G (-68) | appA (Q218*) | oppA (Δ 1bp) | csbC (M1V) | |
| | **EV13-K27** | C | catD (A110V) | srfAB(L496P) | katA-G (-68) | appA (Q218*) | oppA (Δ 1bp) | rpoC (T1143K) | |
| | **EV13-K31** | C | catD (A110V) | srfAB(L496P) | katA-G (-68) | appA (Q218*) | oppA (Δ 1bp) | | |
| | **EV13-K57** | F | ahpF (G495D) | srfAB(Q1900*) | | pckA (R20L) | bdhJ (S333Y) | | |
| | **EV13-K41** | F | ahpF (G495D) | srfAB(Q1900*) | papB(Y155*) | pckA (R20L) | bdhJ (S333Y) | | |
| | **EV13-K82** | E | degU (G224R) | oppA (Δ 1bp) | kinC (K72E) | prkC (A445E) | bcbE (M278I) | ytoQ (D79N) | tlpB (S320G) |
| | **EV13-K83** | D | | | | | | | |
| **PS-218** | **EV218-K2** | A | efeM (E340K) | ΔsrfAB | perR (P22S) | rapA (D194G) | xkdV (S330L) | | |
| | **EV218-K16** | D | skin (Δ 48033bp) | yisY (E254G) | yjcK (A156P) | ylqG (P457Q) | epsE (Δ 1bp) | yrdP (Δ 1bp) | |
| | **EV218-K20** | F | skin (Δ 48033bp) | yisY (E254G) | yjcK (A156P) | ylqG (P457Q) | epsE (Δ 1bp) | yrdP (Δ 1bp) | |
| | **EV218-K27** | F | skin (Δ 48033bp) | yisY (E254G) | yjcK (A156P) | ylqG (P457Q) | epsE (Δ 1bp) | yrdP (Δ 1bp) | |
| | **EV218-K12** | D | hypot. prot. (H135Y) | ahpC (W170R) | resC (A64V) | | | | |
| | **EV218-K56** | F | skin (Δ 48033bp) | yrdP (Δ 1bp) | rhgX (ins. A) | yfkN (A256G) | ylqG (P457Q) | | |
| | **EV218-K57** | E | glnQ (E25Q) | essC (E1425K) | fabZ (D126Y) | cidR (R4H) | yybG (P47S) | | |

### Supplementary Table 2: Table of evolved clones tested for surfactant concentration

*Supplement table 2: Table of clones tested for surfactants production using Drop collapse assay. Clone's contact strain is indicated on the left and their swarming phenotype as a clone's background colour (swarming as white, impaired swarming in light blue and non-swarming in dark blue) with more specifically described swarming phenotype in the 6$^{th}$ column. Abbreviated name indicated in Figure 5 is also provided along with average surfactant concentrations determined according to Drop collapse assay.*

| Contact strain | Population | Clone | Average surfactant concentration (µg/ml) | Swarming phenotype |
|---|---|---|---|---|
| PS-216 | A | EV216-K1 | 101,2 | Non-dendritic swarming |
| | A | EV216-K4 | 114,7 | Non-dendritic swarming |
| | C | EV216-K7 | 134,6 | Non-dendritic swarming |
| | A | EV216-K10 | 140,6 | Swarming |
| | E | EV216-K12 | 102,6 | Swarming |
| | E | EV216-K59 | 96,6 | Swarming |
| | F | EV216-K106 | 96,9 | Swarming |
| | E | EV216-K107 | 104,0 | Swarming |
| | E | EV216-K108 | 121,6 | Swarming |
| | F | EV216-K109 | 117,7 | Swarming |
| | E | EV216-K110 | 101,3 | Swarming |
| | E | EV216-K111 | 112,3 | Swarming |
| | E | EV216-K112 | 126,1 | Swarming |
| | E | EV216-K113 | 115,8 | Swarming |
| | B | EV216-K3 | 4,3 | Non-dendritic swarming |
| | B | EV216-K6 | 8,7 | Non-dendritic swarming |
| | B | EV216-K9 | 3,3 | Non-dendritic swarming |
| | E | EV216-K15 | 183,5 | Swarming |
| | D | EV216-K16 | 2,2 | Non-swarming |
| | E | EV216-K25 | 4,1 | Non-swarming |
| | D | EV216-K36 | 3,3 | Non-swarming |
| | D | EV216-K42 | 3,5 | Non-swarming |
| | D | EV216-K54 | 3,4 | Non-swarming |
| | F | EV216-K89 | 4,7 | Non-swarming |
| | D | EV216-K105 | 2,9 | Non-swarming |
| PS-13 | D | EV13-K60 | 88,4 | Swarming |

| | | | | |
|---|---|---|---|---|
| | D | EV13-K61 | 90,7 | Swarming |
| | D | EV13-K80 | 129,3 | Swarming |
| | D | EV13-K83 | 117,8 | Swarming |
| | D | EV13-K85 | 89,1 | Swarming |
| | D | EV13-K86 | 100,3 | Swarming |
| | D | EV13-K88 | 98,5 | Swarming |
| | D | EV13-K87 | 56,7 | Swarming |
| | F | EV13-K84 | 25,9 | Swarming |
| | F | EV13-K79 | 3,0 | Non-swarming |
| | E | EV13-K82 | 4,0 | Swarming |
| | C | EV13-K1 | 2,3 | Impaired movement |
| | C | EV13-K8 | 2,8 | Impaired movement |
| | C | EV13-K27 | 2,0 | Non-swarming |
| | C | EV13-K15 | 3,1 | Non-swarming |
| | C | EV13-K20 | 2,3 | Impaired movement |
| | C | EV13-K31 | 1,8 | Impaired movement |
| | F | EV13-K41 | 1,7 | Non-swarming |
| | F | EV13-K57 | 3,4 | Non-swarming |
| | F | EV13-K66 | 2,4 | Non-swarming |
| | F | EV13-K81 | 4,0 | Non-swarming |
| PS-218 | A | EV218-K1 | 126,8 | Swarming |
| | D | EV218-K3 | 110,4 | Non-dendritic swarming |
| | D | EV218-K7 | 123,8 | Non-dendritic swarming |
| | D | EV218-K10 | 112,8 | Non-dendritic swarming |
| | D | EV218-K12 | 119,7 | Swarming |
| | F | EV218-K16 | 100,2 | Non-dendritic swarming |
| | F | EV218-K20 | 95,4 | Non-dendritic swarming |
| | F | EV218-K24 | 104,8 | Non-dendritic swarming |
| | F | EV218-K27 | 95,4 | Non-dendritic swarming |
| | F | EV218-K30 | 142,7 | Non-dendritic swarming |
| | F | EV218-K32 | 195,1 | Swarming |
| | D | EV218-K56 | 114,3 | Swarming |
| | E | EV218-K57 | 81,3 | Swarming |
| | D | EV218-K58 | 119,1 | Swarming |
| | D | EV218-K60 | 129,0 | Swarming |
| | D | EV218-K61 | 117,3 | Swarming |
| | D | EV218-K62 | 86,1 | Swarming |
| | D | EV218-K63 | 93,4 | Swarming |
| | E | EV218-K59 | 61,7 | Swarming |
| | A | EV218-K2 | 11,7 | Impaired movement |
